# The *disconnect2* mutation disrupts the *tjp1b* gene that is required for electrical synapse formation

**DOI:** 10.1101/2022.06.27.497839

**Authors:** Jennifer Carlisle Michel, Abagael M. Lasseigne, Audrey J. Marsh, Adam C. Miller

## Abstract

To investigate electrical synapse formation *in vivo* we used forward genetics to disrupt genes affecting Mauthner cell electrical synapses in larval zebrafish. We identify the *disconnect2* (*dis2*) mutation for its failure to localize neural gap junction channels at electrical synapses. We mapped this mutation to chromosome 25 and identified a splice-altering mutation in the *tjp1b* gene. We demonstrated that the *dis2* mutation disrupts *tjp1b* function using complementation analysis with CRISPR generated mutants. We conclude that the *dis2* mutation disrupts the *tjp1b* gene that is required for electrical synapse formation.

**Description:** Neural networks are circuits of neurons wired together during development that provide an animal with specialized behavioral outputs. Dedicated adhesions called neuronal synapses create sites of communication between neurons and can be categorized as either electrical or chemical. This work identifies a new mutation in *tjp1b* that is shown to be required for electrical synapse formation.

## Manuscript

Vertebrate electrical synapses are gap junction (GJ) channels formed between neurons when two Connexin hemichannels dock (Söhl et al. 2005), creating a direct interneuronal path for ionic and metabolic coupling. An individual electrical synapse contains tens to thousands of GJ channels, which are organized into so-called plaques and have stereotyped morphologies dependent upon location (Nagy et al. 2018). The localization of Connexin proteins to the electrical synapse is thought to be regulated by a network of molecular interactions between the Connexins and intracellular scaffolds (Nagy et al. 2018; Martin et al. 2020). Emerging evidence suggests that complex multimolecular structures regulate electrical synapse formation and function at vertebrate neuronal GJs (Miller et al. 2015; Marsh et al. 2017; Lasseigne et al. 2021), yet the gene identities and functions of these molecules are still poorly understood.

To investigate genes required for electrical synapse formation *in vivo* we used the electrical synapses of the Mauthner cell in larval zebrafish, *Danio rerio* (Fig. 1A). Mauthner cell somas and dendrites are located in the hindbrain where they receive multi-modal sensory input. Each Mauthner sends an axon down the length of the spinal cord where they coordinate a fast escape response to threatening stimuli (Eaton et al. 1977; Jacoby and Kimmel 1982; Liu and Fetcho 1999). Our analysis focused on the ‘club ending’ (CE) synapses formed between auditory afferents of the eighth cranial nerve and the Mauthner cell’s lateral dendrite (Yao et al. 2014) and *en passant* electrical synapses between the Mauthner cell axon and Commissural Local (CoLo) interneurons (Satou et al. 2009). The Mauthner and CoLo neurons can be visualized using the transgenic line *zf206Et(Tol-056)*, which expresses green fluorescent protein (GFP) in both neuron types (Satou et al. 2009). The electrical synapses of the Mauthner circuit are heterotypic; that is, hemichannels form from unique Connexin proteins on each side of the synapse. Cx35.5, encoded by the gene *gap junction delta 2a* (*gjd2a*), is used exclusively presynaptically, while Cx34.1 (*gjd1a*) is used exclusively postsynaptically (Miller et al. 2017). Both Connexins can be visualized by immunostaining using a polyclonal antibody against the human Cx36 protein (Fig. 1B,C). We performed a forward genetic screen using N-ethyl-N-nitrosourea (ENU) to generate random mutations and identified the *disconnect2* (*dis2*) mutation that caused a loss of detectable Cx36 staining at both the CE and M/CoLo synapses with no apparent effect on neuronal morphology (Fig. 1D-G). These results support the notion that the *dis2* mutation affects a gene required for electrical synapse formation.

**Figure 1.**
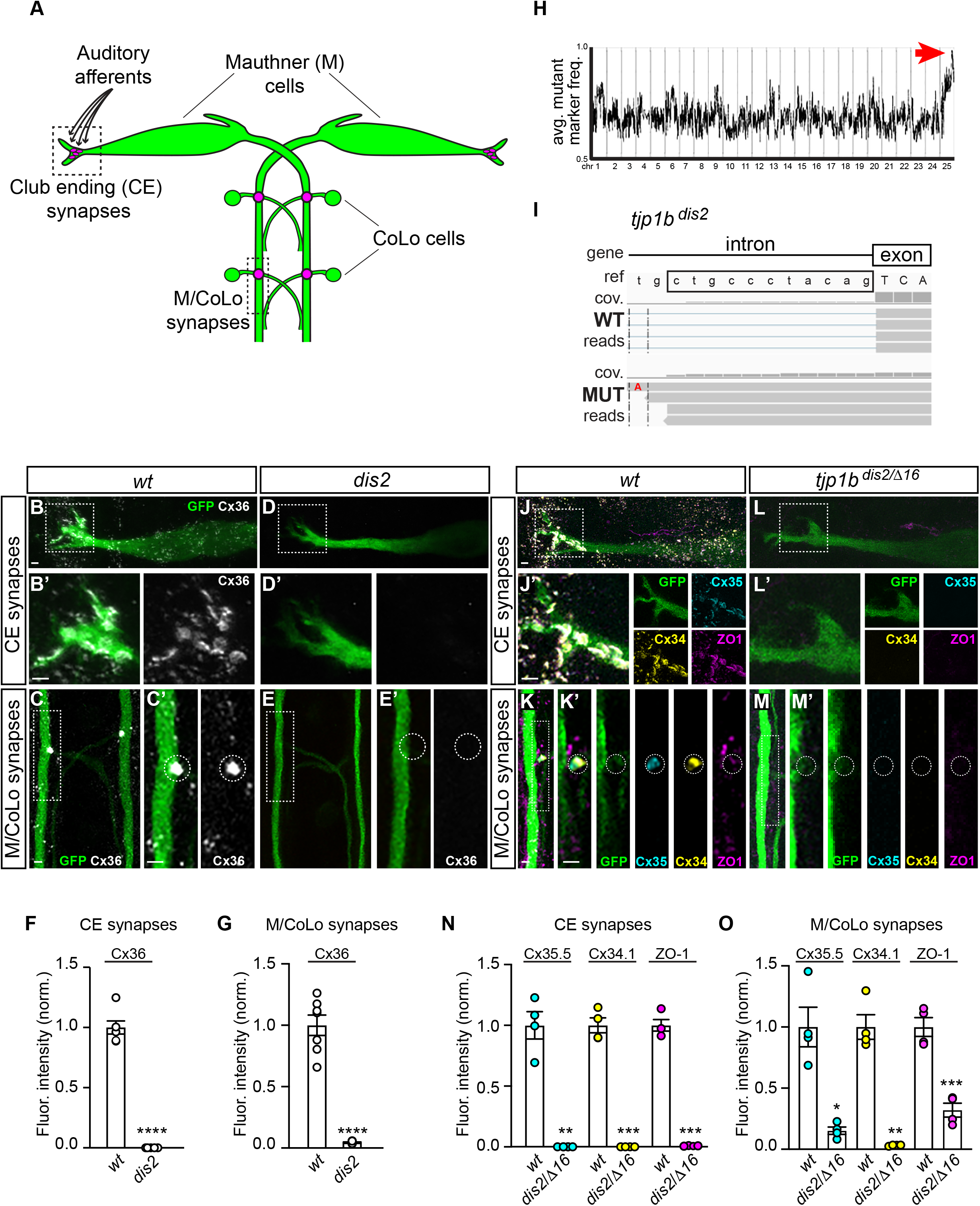
The *disconnect2* mutation disrupts the *tjp1b* gene required for electrical synapse formation. **A:** Diagram of the Mauthner cell circuit illustrating the electrical synapses of interest. The image represents a dorsal view with anterior on top. Boxed regions indicate the stereotypical synaptic contacts used for analysis. Presynaptic auditory afferents contact the postsynaptic Mauthner cell lateral dendrite in the hindbrain forming Club Ending (CE) synapses. Presynaptic Mauthner axons form *en passant* electrical synapses with the postsynaptic CoLo interneurons (M/CoLo synapses) in the spinal cord (2 of 30 segments are depicted). **B–E:** Confocal images of Mauthner circuit neurons and stereotypical electrical synaptic contacts in 5 dpf *zf206Et* zebrafish larvae from *wildtype* (*wt*) (**B, C**) and *dis2*^*-/-*^ mutant (**D, E**) animals. Animals are stained with anti-GFP (green) and anti-Cx36 (white). Scale bars = 2 *µ*m. Boxed regions denote stereotyped location of electrical synapses and regions are enlarged in neighboring panels. **B, D**: Images of the Mauthner cell body and lateral dendrite in the hindbrain. Images are maximum intensity projections of ∼10 *µ*m. In B’ and D’, images are maximum-intensity projections of ∼5 *µ*m and neighboring panel shows the Cx36 channel. **C, E**: Images of the sites of contact of Mauthner/CoLo processes in the spinal cord. Images are maximum-intensity projections of ∼4 *µ*m. In C’ and E’, the white dashed circle denotes the M/CoLo site of contact and the neighboring panel shows the Cx36 channel. **F, G:** Quantification of Cx36 fluorescence intensities at CE (**F**) and M/CoLo (**G**) synapses for the noted genotypes. The height of the bar represents the mean of the sampled data normalized to the *wt* average. Circles represent the normalized value of each individual animal. All CEs (∼10) of both Mauthner cells were sampled per animal, and between 12 and 18 M/CoLo synapses were sampled per animal. CE synapses: *wt* n=6, *dis2*^*-/-*^ n=5, p<0.0001; M/CoLo synapses: *wt* n=7, *dis2*^*-/-*^ n=7, p<0.0001. Error bars are ± SEM. **H:** Genome wide RNA-seq-based mapping data. The average frequency of mutant markers (black marks) is plotted against genomic position. A single region on chromosome 25 was linked to the *dis2* mutation (red arrow). **I:** Mutant reads are shown aligned to the reference genome identifying a T to A transversion (highlighted in red font) creating a premature splice acceptor site in the intron and introducing 11 base pairs of intronic sequence (boxed in black) into the transcript that causes a frameshift. A sample of aligned reads are shown as grey boxes. The coverage (cov.) of aligned reads is depicted as a histogram at each genomic position. **J–M:** Confocal images of Mauthner circuit neurons and stereotypical electrical synaptic contacts in 5 dpf *zf206Et* zebrafish larvae from *wt* (**J, K**) and *tjp1b*^*dis2/Δ16*^ (**L, M**) animals. Animals are stained with anti-GFP (green), anti-Cx35.5 (cyan), anti-Cx34.1 (yellow), and anti-ZO1 (magenta). Scale bars = 2 *µ*m. Boxed regions denote stereotyped location of electrical synapses and regions are enlarged in neighboring panels. **J, L**: Images of the Mauthner cell body and lateral dendrite in the hindbrain. Images are maximum intensity projections of ∼20 *µ*m. In J’ and L’, images are maximum-intensity projections of ∼10 *µ*m and neighboring panels show the individual channels. **K, M**: Images of the sites of contact of M/CoLo processes in the spinal cord. Images are maximum-intensity projections of ∼8 *µ*m. In K’ and M’, images are from a single 0.4 *µ*m Z-plane and the white dashed circle denotes the location of the M/CoLo site of contact. Neighboring panels show individual channels. **N, O:** Quantification of Cx35.5 (cyan), Cx34.1 (yellow), and ZO1 (magenta) fluorescence intensities at CE (**N**) and M/CoLo (**O**) synapses for the noted genotypes. The height of the bar represents the mean of the sampled data normalized to the *wt* average. Circles represent the normalized value of each individual animal. CE synapses: *wt* n=4, *tjp1b*^*dis2/Δ16*^ n=4, Cx35.5 p=0.003, Cx34.1 p=0.0005, ZO1 p=0.0002; M/CoLo synapses: *wt* n=4, *tjp1b*^*dis2/Δ16*^ n=4, Cx35.5 p=0.0121, Cx34.1 p=0.0024, ZO1 p=0.0005. Error bars are ± SEM.

To identify the gene affected by the *dis2* mutation we used an RNA-sequencing-based approach (Miller et al. 2013) and mapped the mutation to an ∼1.5 megabase region on chromosome 25 (Fig. 1H). Using the RNA-seq data within this region we found that mutants harbored a single nucleotide polymorphism (SNP) that introduced a novel splice acceptor within the *tight junction 1b (tjp1b)* gene. This introduced 11 base pairs (bps) of what is normally intronic sequence into the transcript. The additional nucleotides are inserted at position 1352 of the 7626bp transcript (ENSDART00000155992.3) causing a frameshift in the remaining sequence (Fig. 1I). In previous work we used a CRISPR-based reverse genetic screen and identified the *tjp1b* gene, which encodes the cytoplasmic scaffolding protein ZO1b, as being required for electrical synapse formation (Shah et al. 2015; Marsh et al. 2017). Therefore, we tested whether the *dis2* mutation could complement our CRISPR generated, 16 bp deletion (*tjp1b*^*Δ16*^). We found that trans-heterozygote *dis2 / tjp1b*^*Δ16*^ animals failed to localize both presynaptic Cx35.5 and postsynaptic Cx34.1 to Mauthner electrical synapses (Fig. 1J-O). Moreover, we found that ZO1 staining, which recognizes the protein encoded by the *tjp1b* gene that normally co-localizes with Connexin staining at Mauthner electrical synapses, was greatly reduced in the *dis2 / tjp1b*^*Δ16*^ animals (Fig. 1J-O). This phenocopies the results we observe in homozygous *tjp1b*^*Δ16/Δ16*^ mutant animals (Lasseigne et al. 2021). We renamed the *dis2* mutation *tjp1b*^*b1435*^ and conclude that the mutation disrupts the *tjp1b* gene, which is required for electrical synapse formation.

Here we identify a new mutant allele of *tjp1b*, and we show it is required for electrical synapse formation. Our previous work has shown that the ZO1b protein, encoded by the *tjp1b* gene, is exclusively localized postsynaptically at Mauthner electrical synapses, where it biochemically interacts with the postsynaptic Cx34.1, an interaction required for the structure and function of neuronal GJs (Lasseigne et al. 2021). Growing evidence suggests that electrical synapses are complex and asymmetric structures, analogous to their chemical synapse cousins (Martin et al. 2020). Identifying new mutant alleles of the genes involved in neuronal GJ formation will provide critical tools to uncover the complexity of electrical synapses *in vivo*.

## Methods

### Zebrafish

Zebrafish, *Danio rerio*, were bred and maintained in the University of Oregon fish facility at 28°C on a 14 hr on and 10 hr off light cycle with approval from the Institutional Animal Care and Use Committee. Animals were staged using standard procedures (Kimmel et al. 1995). The *dis2* allele (*tjp1b*^*b1435*^) was isolated from an early-pressure, gynogenetic diploid screen (Walker et al. 2009) using ENU as a mutagen and was maintained in the *zf206Et(Tol-056)* background (Satou et al. 2009). The *tjp1b*^*b1370*^ mutant line contains a 16 bp deletion in the tjp1b gene (Marsh et al. 2017). Mutant lines were genotyped for all experiments, and all immunohistochemistry was performed at 5 days post fertilization (dpf).

### RNA-seq-based mutant mapping

Total RNA was extracted from *dis2* mutants and *wildtype* siblings, and cDNA libraries were created using standard Illumina TruSeq protocols. Each library was individually barcoded allowing for identification after multiplexed sequencing on an Illumina HiSeq 2000 machine. There were ∼60 million reads per pool and these were aligned to the zebrafish genome (Zv9.63) using TopHat/Bowtie, an intron and splice aware aligner (Trapnell et al. 2012). Single nucleotide polymorphisms (SNPs) were identified using the SAMtools mpileup and bcftools variant caller (Li et al. 2009). Custom R scripts (Miller et al. 2013) were used to identify high quality mapping SNPs in the wildtype pool and these positions were then assessed in the mutant pool for their frequency. The average allele frequency, using a sliding-window of 50-neighboring loci, was plotted across the genome and linkage was identified as the region of highest average frequency. Within the linked region, candidate mutations were identified using a combination of custom R scripts and existing software (Variant Effect Predictor (McLaren et al. 2010), Cufflinks (Trapnell et al. 2012)). Details can be found at www.RNAmapper.org (Miller et al. 2013).

### Immunohistochemistry and confocal imaging

Anesthetized, 5 dpf larvae were fixed for 3 hours in 2% trichloroacetic acid in PBS. Fixed tissue was washed in PBS plus 0.5% Triton X-100, followed by standard blocking and antibody incubations (Martin et al. 2022). Primary antibody mixes included combinations of the following: rabbit anti-Cx36 (Invitrogen, 36-4600, 1:200), chicken anti-GFP (Abcam, ab13970, 1:500), rabbit anti-Cx35.5 (Fred Hutch Antibody Technology Facility, clone 12H5, 1:800), mouse IgG2A anti-Cx34.1 (Fred Hutch Antibody Technology Facility, clone 5C10A, 1:350), and mouse IgG1 anti-ZO1 (Invitrogen, 33–9100, 1:350). All secondary antibodies were raised in goat (Invitrogen, conjugated with Alexa-405,–488, −555, 594, or −633 fluorophores, 1:500). Tissue was then cleared stepwise in a 25%, 50%, 75% glycerol series, dissected, and mounted in ProLong Gold antifade reagent (ThermoFisher). Images were acquired on a Leica SP8 Confocal using a 405-diode laser and a white light laser set to 499, 553, 598, and 631 nm, depending on the fluorescent dye imaged. Each laser line’s data was collected sequentially using custom detection filters based on the dye. Quantitative images of the Club Endings (CEs) were collected using a 63x, 1.40 numerical aperture (NA), oil immersion lens, and images of M/Colo synapses were collected using a 40x, 1.20 NA, water immersion lens. For each set of images, the optimal optical section thickness was used as calculated by the Leica software based on the pinhole, emission wavelengths, and NA of the lens. Within each experiment where fluorescence intensity was to be quantified, all animals were stained together with the same antibody mix, processed at the same time, and all confocal settings (laser power, scan speed, gain, offset, objective, and zoom) were identical. Multiple animals per genotype were analyzed to account for biological variation. To account for technical variation, fluorescence intensity values for each region of each animal were an average across multiple synapses.

### Analysis of confocal imaging

For fluorescence intensity quantitation, confocal images were processed and analyzed using FiJi software (Schindelin et al. 2012). To quantify staining at M/Colo synapses, a standard region of interest (ROI) surrounding each M/CoLo site of contact was drawn and the mean fluorescence intensity was measured. For the quantification of staining at the club endings, confocal z-stacks of the Mauthner soma and lateral dendrite were cropped to 36.08 *µ*m x 36.08 *µ*m centered around the lateral dendritic bifurcation. Using the SciPy (Virtanen et al. 2020) and scikit-image (van der Walt et al. 2014) computing packages, the cropped stack was then cleared outside of the Mauthner cell, a 3^3^ median filter was applied to reduce noise, and a standard threshold was set within each experiment to remove background staining. The image was then transformed into a max intensity projection and the integrated density of each stain within the Mauthner cell was extracted. Figure images were created using FiJi, Photoshop (Adobe), and Illustrator (Adobe).

### Statistical analysis

Statistical analyses were performed using Prism software (GraphPad). For all experiments, values were normalized to *wildtype* control animals, and n represents the number of fish used. An unpaired t-test with Welch’s correction was performed and error bars represent standard error of the mean.

## Author contributions

Jennifer Carlisle Michel: Formal analysis, Methodology, Project administration, Visualization, Writing - original draft, Writing - review and editing

Abagael M. Lasseigne: Formal analysis, Investigation, Methodology, Validation

Audrey M. Marsh: Investigation, Methodology, Validation

Adam C. Miller: Conceptualization, Data curation, Formal analysis, Funding acquisition, Investigation, Methodology, Project administration, Resources, Supervision, Visualization, Writing – original draft, Writing - review and editing

## Acknowledgments

We thank the University of Oregon Fish Facility for superb animal care. We appreciate the discussions and advice from all lab members and from members of the Institute of Neuroscience and the Institute of Molecular Biology at the University of Oregon.

## Reagents

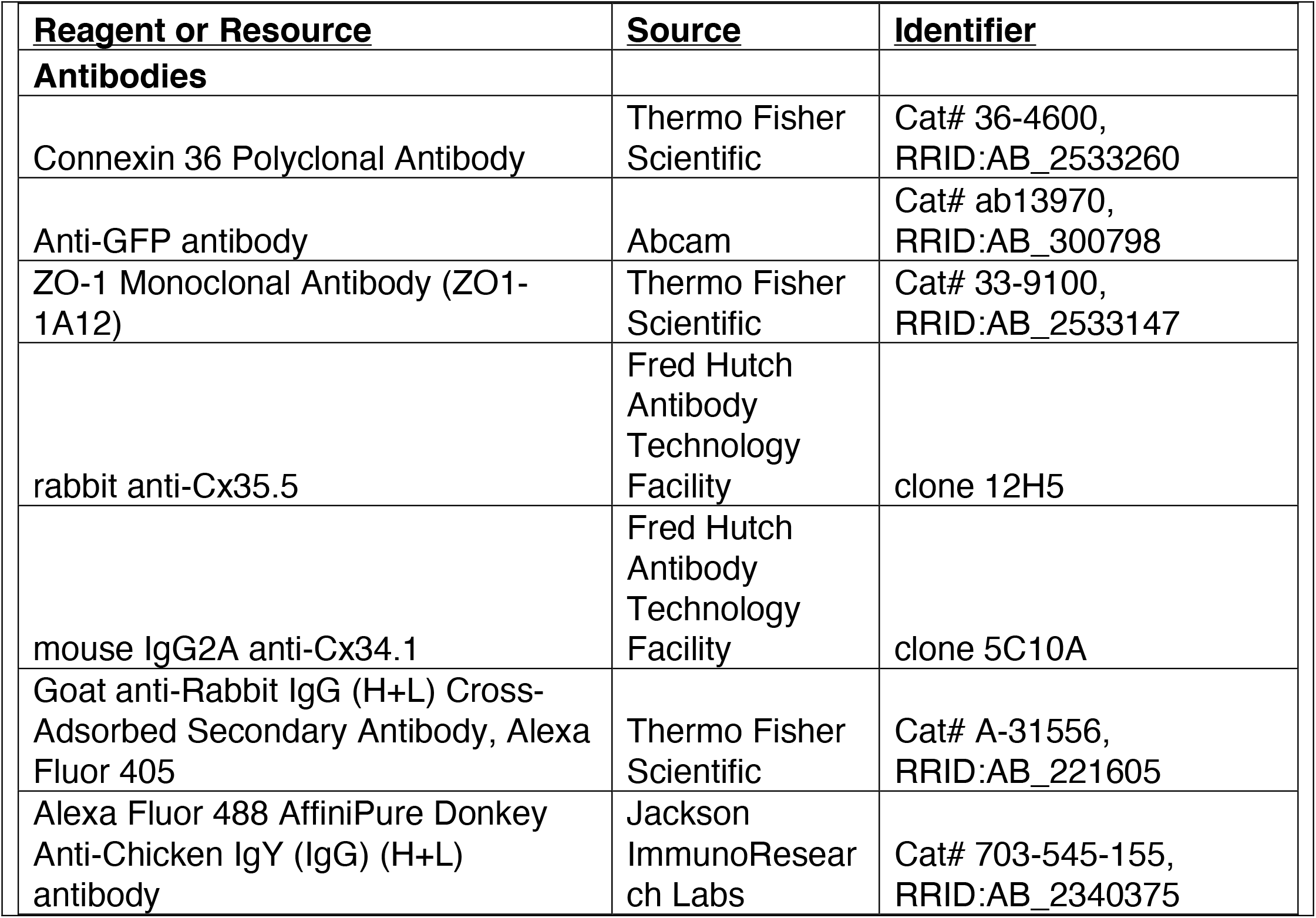
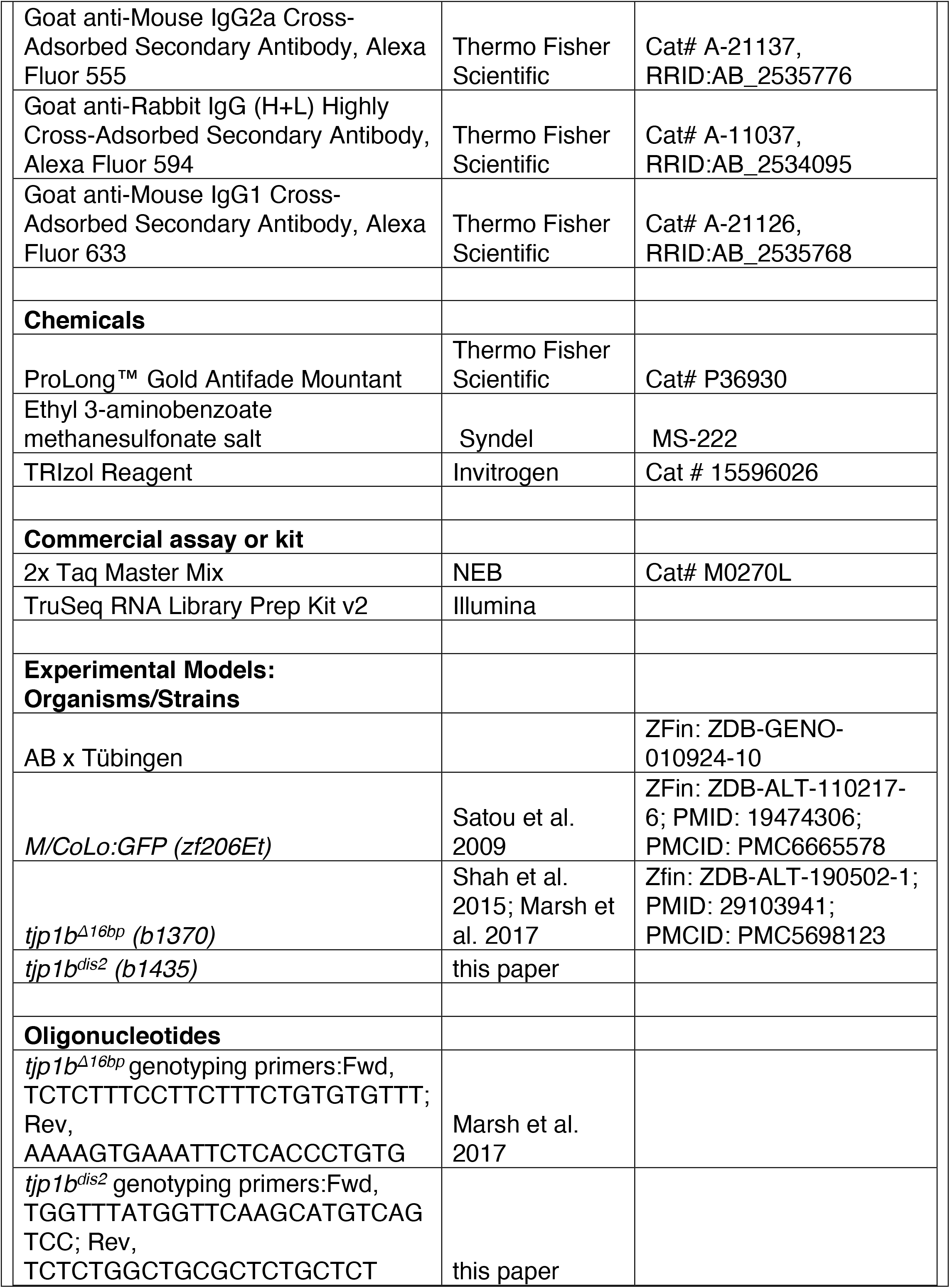
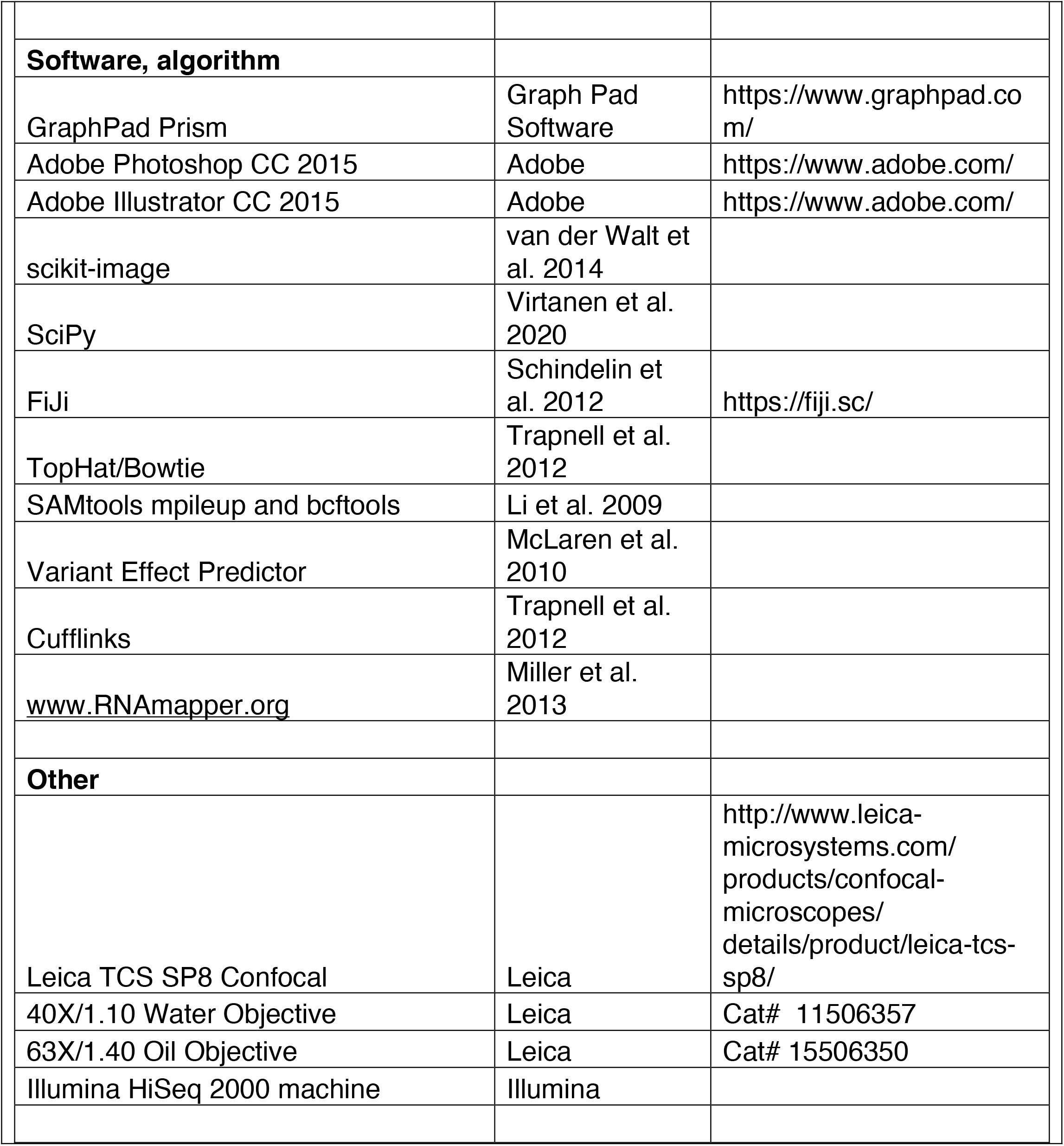

